# Development of SacB-based Counterselection for Efficient Allelic Exchange in *Fusobacterium nucleatum*

**DOI:** 10.1101/2024.08.16.608263

**Authors:** Peng Zhou, G C Bibek, Bo Hu, Chenggang Wu

## Abstract

*Fusobacterium nucleatum*, prevalent in the oral cavity, is significantly linked to overall human health. Our molecular comprehension of its role in oral biofilm formation and its interactions with the host under various pathological circumstances has seen considerable advancements in recent years, primarily due to the development of various genetic tools for DNA manipulation in this bacterium. Of these, counterselection-based unmarked in-frame mutation methods have proved notably effective. Under suitable growth conditions, cells carrying a counterselectable gene die, enabling efficient selection of rare defined allelic exchange mutants. The *sacB* gene from *Bacillus subtilis*, encoding levansucrase, is a widely used counterselective marker partly due to the easy availability of sucrose. Yet, its potential application in *F. nucleatum* genetic study remains untested. We demonstrated that *F. nucleatum* cells expressing *sacB* in either a shuttle or suicide plasmid exhibit a lethal sensitivity to supplemental sucrose. Utilizing sucrose counterselection, we created an in-frame deletion of the *F. nucleatum tonB* gene, a critical gene for energy-dependent transport processes in Gram-negative bacteria, and a precise knockin of the luciferase gene immediately following the stop codon of the *hslO* gene, the last gene of a five-gene operon possible related to the natural competence of *F. nucleatum*. Post counterselection with 5% sucrose, chromosomal plasmid loss occurred in all colonies, leading to gene alternations in half of the screened isolates. This *sacB*-based counterselection technique provides a reliable method for isolating unmarked gene mutations in wild-type *F. nucleatum*, enriching the toolkit for fusobacterial research.

**IMPORTANCE:** Investigations into *Fusobacterium nucleatum*’s role in related diseases significantly benefit from the strategies of creating unmarked gene mutations, which hinge on using a counterselective marker. Previously, the *galk*-based allelic exchange method, while effective, faced an inherent limitation – the need for a modified host. This study aims to surmount this limitation by substituting *galK* with *sacB* for gene modification in *F. nucleatum*. Our application of the *sacB*-based methodology successfully yielded a *tonB* in-frame deletion mutant and a luciferase gene knockin at the precise chromosomal location in the wild-type background. The new method augments the existing toolkit for *F. nucleatum* research and has far-reaching implications due to the easy accessibility to the counterselection compound sucrose. We anticipate its broader adoption in further exploration, thereby reinforcing its critical role in propelling our understanding of *F. nucleatum*.

## INTRODUCTION

In the human subgingival biofilm, a richly diverse microbial community hosting over 400 microbial species, *Fusobacterium nucleatum*, an anaerobic, Gram-negative bacterium, thrives prominently (1–3). Regardless of the health status of dental plaque, *F. nucleatum* maintains the second most abundant position within this biofilm; this presence significantly escalates amidst periodontal inflammation (4, 5). Beyond its association with periodontitis, evidence suggests that when disseminated to extra-oral sites, *F. nucleatum* could potentially influence a host of systemic diseases, including cardiovascular disease, adverse pregnancy outcomes, rheumatoid arthritis, and various cancer types (6, 7).

The clinical importance of *F. nucleatum* has spurred extensive research in the last twenty years (6, 7). Advances in our understanding of its biology and role in diseases are mainly due to genetic tools enabling DNA manipulation, particularly ones that create defined gene mutations. Two such mutation methods are insertion duplication mutagenesis (8, 9) and marked allelic replacement mutagenesis (10, 11). The former method employs a plasmid carrying part of the desired gene and a thiamphenicol-resistance gene (*catP*) as a selectable marker. Once introduced into fusobacterial cells, this plasmid is integrated into the chromosome via homologous recombination, disrupting the target gene and producing a truncated protein. Prominent mutations resulting from this method include those in *fap2* (8), *radD* (*12*), *cmpA* (*13*), and *FplA* (14) genes. Despite its directness, this method presents several limitations. It is ineffective for small genes and cannot investigate the functionality of genes within a multi-gene operon that is not the last gene due to the potential polar effect it may induce. However, marked allelic replacement mutagenesis can partly address these limitations. This strategy involves a mutagenesis cassette carrying an antibiotic resistance marker (*catP* or *ermF-ermAM* for clindamycin) and upstream and downstream homology arms that align with the chromosome region adjacent to the target gene. Notable mutations from this method include those in *recA* (10), *fadA* (11), and *aid1*(15) genes. Although practical, these methods have restrictions in creating nuanced mutations, and the lack of selectable markers limited the possible mutations per strain. Unmarked allelic exchange, however, can circumvent these obstacles.

The unmarked allelic exchange usually involves two steps. In the first step, two PCR-amplified fragments, which are similar in size and homologous to sequences adjacent to the mutation site, are generated. These fragments are then positioned side by side within a suicide vector, forming an in-frame deletion construct. Integration of this construct into a cell’s chromosome occurs through a single crossover event at one of the homologous fragments. Positive selection, such as antibiotic resistance, is employed to choose integrated clones. In the second step, the integrated clones are allowed to propagate in the absence of antibiotics, facilitating a second crossover between the duplicated homologous regions resulting from insertion (16). This double crossover leads to plasmid loss from the cells. Since the suicide vector contains two similar-sized homologous fragments, the secondary recombination has an equal probability of occurring at either site. As a result, the outcome can be either an in-frame deletion mutation or a reversion back to the wild-type genotype, both with an equal likelihood (Figure 1A). However, the screening process for these recombinants poses a challenge due to the infrequent occurrence of the double crossover event. It involves the laborious task of manually identifying hundreds or even thousands of individual colonies. Thus, counterselection strategies are often employed to make this screening procedure more efficient and streamlined.

**Figure 1.**
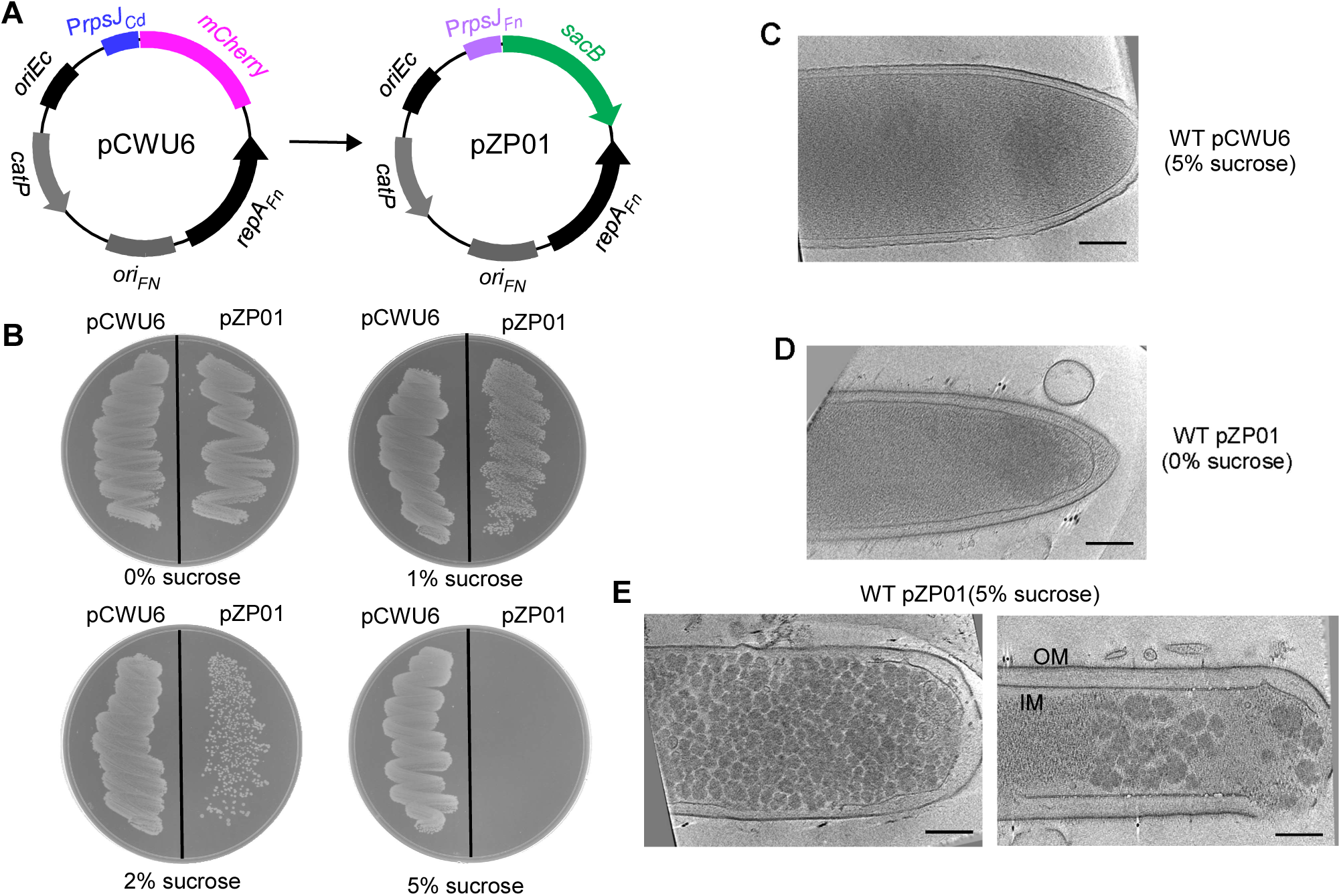
Sucrose Tolerance Assays for SacB-Expressing *F. nucleatum* Strain. (A) Construction of the *E. coli/F. nucleatum* shuttle vector pPZ01. The *sacB* gene was expressed under the control of the *F. nucleatum rpsJ* promoter. (B) Sucrose tolerance assays. *F. nucleatum* strains carrying either the pPZ01 or pCWU6 plasmids were streaked onto TSPC agar plates supplemented with 0%, 1%, 2%, or 5% sucrose and incubated anaerobically at 37°C for 48 hours. The experiments were repeated three times, with a representative result shown here. (C, D, E) Cryo-electron microscopy images illustrate the effects of SacB expression on cells in the presence and absence of sucrose. IM: inner membrane; OM: outer membrane. Scale bars represent 50 nm.

Applying a counterselective strategy necessitates a counterselectable gene marker housed within an in-frame deletion plasmid construct for negative selection. In this context, the gene marker product triggers self-destruction of the cell containing it. As a result, transformed cells that have integrated a suicide vector bearing a counterselectable marker through a single crossover event will retain a chromosomal copy of the marker gene. Subsequently, these cells are eradicated when exposed to either the counterselective compound or the inducer (when the product of the counterselection gene is intrinsically toxic). This implies that only strains that have completed the second crossover to remove the plasmid will remain viable. In this scenario, approximately half of the emerging colonies should exhibit an in-frame deletion, while the rest are expected to be of the wild type. The *galK* gene, often used as a counterselection marker, encodes galactokinase, a key enzyme in galactose metabolism (16). This enzyme converts D-galactose into galactose-1-phosphate, and it can also transform 2-dexoygalactose-1-phosphate into a non-metabolizable and toxic substance, 2-deoxygalactose-1-phosphate. Our laboratory, among others, has effectively used *galK* as a counterselection marker in *F. nucleatum*, generating some in-frame deletions (17–20). However, the *galk*-based system has limitations since *F. nucleatum* naturally carries a copy of the *galK* gene. To employ *galK* for counterselection, we need to create a *galK*-minus recipient strain (17). Although the *galK* gene doesn’t interfere with the phenotypes under our study, primarily those concerning RadD-mediated fusobacterial coaggregation *in vitro* (18), its deletion might influence other aspects of this bacterium’s virulence and physiology. This is due to the *galK*’s function linked to galactose utilization. So, the deletion of *galk* affects the galactose level in *F. nucleatum*’s growth environment. Current studies revealed galactose modulates fusobacterial biofilm formation by inhibiting autoinducer 2, a quorum-sensing molecular (21). In addition, galactose can alter the function of Fap2, a crucial adhesin that binds to galactose, contributing to cell-cell coaggregation and host adhesion (22, 23). Furthermore, *F. nucleatum* rapidly uptakes galactose to manufacture sugar polymers, a strategy that likely aids its survival in the nutrient-poor conditions of the subgingival pocket (24). More importantly, UDP-galactose, the intermediate product of galactose utilization, is vital for preserving the cell envelope and producing exopolysaccharides, key in biofilm formation as seen in certain bacteria (25, 26). Consequently, the fusobacterial mutants generated from the *galK*-based system might not be optimal for every fusobacterial study, particularly those involving animal infection models. Therefore, the need for an alternative counterselection marker to replace *galK* in *F. nucleatum* genetic studies is pressing.

To meet this need, we have assessed several prevalent counterselection markers, including *pheS*, *I-sceI*, *sacB*, and toxic genes such as *mazF* and *hicA*. Our investigations revealed that *sacB* and *hicA* (27) effectively serve counterselection markers in the wild-type context of *F. nucleatum*. SacB, a gene initially isolated from *Bacillus subtilis* (27), encodes levansucrase, a secreted enzyme synthesizing levan from sucrose, a process harmless to *B. subtilis*. However, when *sacB* is expressed in other bacteria, it provokes a deadly sensitivity to sucrose (28, 29). The underlying theory for this sucrose toxicity is that the buildup of levan in the cytoplasm or periplasm can disrupt the normal physiological functions of cells, thereby leading to cell lysis (16, 30). Given this mechanism, the *sacB* gene has broad application as a counter-selection marker, particularly in gene deletion procedures that utilize a double-crossover event. This has been successfully implemented in a wide range of both Gram-negative and Gram-positive bacteria (16, 30). Notably, however, such a technique has yet to be described in *F. nucleatum*

In this study, we report the development of a *sacB*-based in-frame mutation system in *F. nucleatum*, the blueprint of which is depicted in Figure 2A. Using this system, we created a markerless in-frame deletion in the *F. nucleatum tonB* gene and an in-frame insertion of a luciferase gene after the stop codon of the *hslO* gene. Counterselection led to consistent plasmid loss in all tested colonies, and about half of the screened isolated grown on 5% sucrose plates were in-frame mutation strains. This *sacB*-based counterselection technique offers a reliable means to generate unmarked gene mutations in the wild-type background of *F*. *nucleatum*, thereby expanding the suite of genetic tools available for fusobacterial research.

**Figure. 2.**
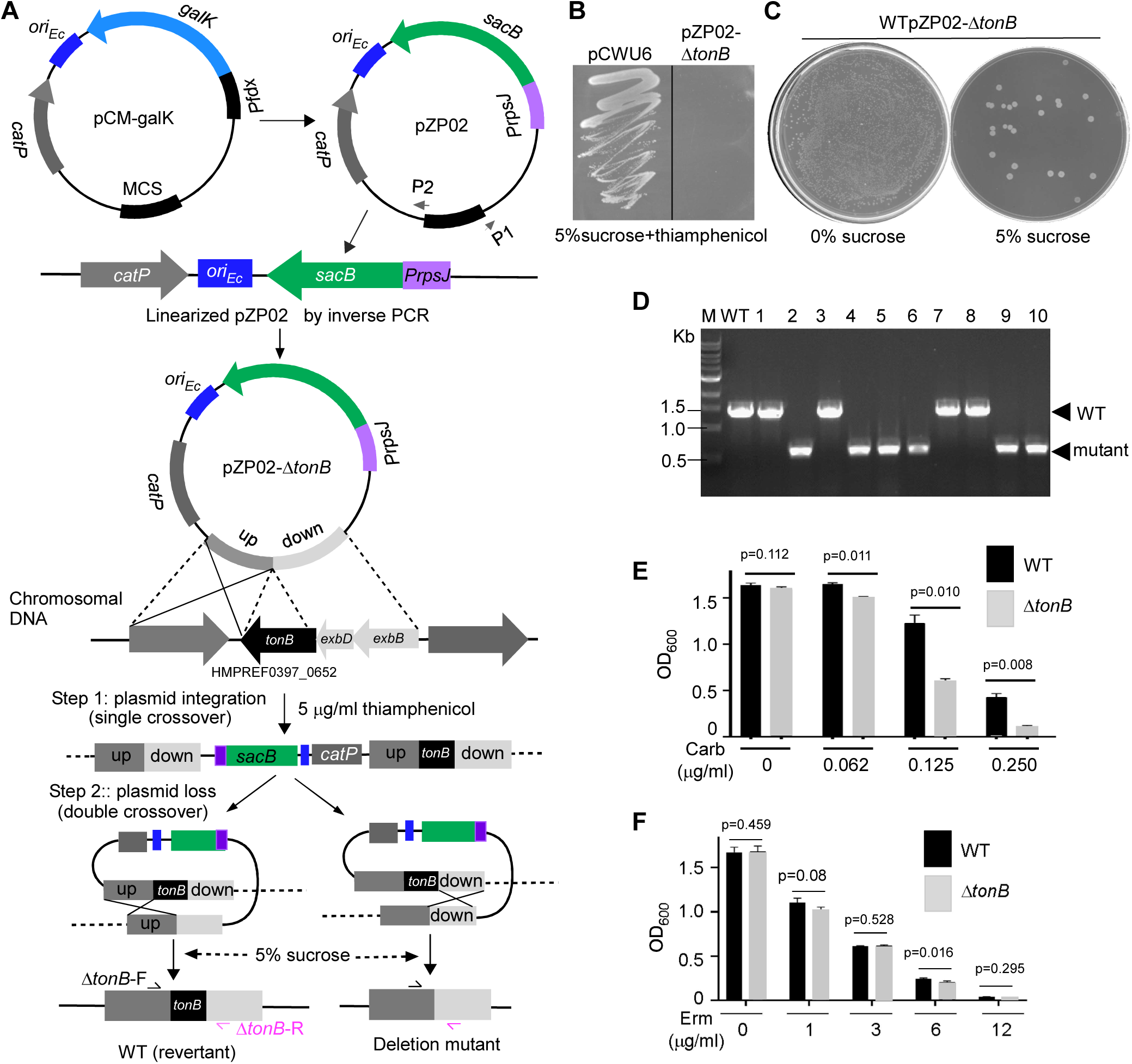
Using *sacB* as a counter-selective marker for *tonB* gene deletion in F. nucleatum. (A) Workflow for *tonB* gene deletion using the pPZ02 plasmid, expressing *sacB*. The plasmid was derived from pCM-galK. Linearization of pPZ02 was achieved by inverted PCR with primers P1/P2, followed by PCR amplification and fusion of 1.5 kb fragments of the *tonB* gene’s flanking regions. The fused fragment was ligated into linearized pPZ02 using Gibson assembly, creating pPZ02-ΔtonB. Gene deletion in *F. nucleatum* was achieved through a two-step allelic exchange: (1) Selection with 5 µg/mL thiamphenicol and (2) Counter-selection with 5% sucrose. (B) Confirmation of sucrose sensitivity due to *sacB* expression. The pPZ02-Δ*tonB* integrated strain was grown in the presence of sucrose and antibiotics for 3 days at 37°C, with wild-type *F. nucleatum* containing pCWU6 as a control. (C) Sucrose counter-selection results for *tonB* deletion. A 1,000-fold diluted overnight culture of the pPZ02-ΔtonB strain was plated on TSPC agar with 0% or 5% sucrose to isolate plasmid-excised cells. (D) *tonB* deletion verification by colony PCR using primers ΔtonB-F/ΔtonB-R. The wild-type yields a 1.5 kb amplicon, while the deletion mutant yields a 0.5 kb product. Six out of ten colonies were confirmed as deletion mutants. (E) Carbenicillin resistance assay. Wild-type and Δ*tonB* strains were treated with 0 to 0.25 µg/mL carbenicillin at 37°C for 24 hours in an anaerobic chamber. (F) Erythromycin sensitivity test. Wild-type and Δ*tonB* strains were incubated with 0 to 12 µg/mL erythromycin at 37°C for 24 hours in an anaerobic chamber. Bacterial growth was measured by OD_600_ for (E) and (F). Results are from three independent experiments, presented as means ± SD.

## RESULTS

### Expression of *sacB* in *F. nucleatum* confers sucrose toxicity

To assess the utility of *sacB* as a counterselection marker in *F. nucleatum*, we created the plasmid pZP01 (Fig. 1A), derived from the shuttle plasmid pCWU6 in *E. coli/F. nucleatum* systems (17). In pZP01, the *sacB* expression was controlled by the *rpsJ* promoter, a strong and constitutively active promoter from *F. nucleatum* (17). Following the transformation of pZP01 into wild-type cells of *F. nucleatum*, we investigated the sensitivity of the resulting transformants to sucrose by conducting growth assays on agar plates supplemented with varying sucrose concentrations.

As demonstrated in Figure 1B, the presence of 1% sucrose did not hinder cell growth, while 2% sucrose led to partial growth inhibition. Remarkably, at a sucrose concentration of 5%, cells expressing *sacB* exhibited complete loss of viability. In contrast, the control group of cells carrying the empty plasmid pCWU6 remained unaffected by the presence of sucrose (Fig. 1B).

To gain deeper insight into the underlying mechanism responsible for the lethal effect of *sacB* in the presence of 5% sucrose, we employed cryo-electron microscopy to examine cells containing pZP01 or pCWU6. Notably, cells expressing *sacB* and cultivated with 5% sucrose displayed a remarkable accumulation of granular sugar polymers throughout the cytoplasm (Fig. 1E). Furthermore, these cells exhibited distinct morphological alternations, deviating from the characteristic spindle-like tips typically observed in normal fusobacterial cells. Ultimately, these architectural changes resulted in cell rupture (Fig.1E). Conversely, the absence of sucrose abolished the formation of granular sugar polymers (Fig.1D). Similarly, cells carrying the empty plasmid pCWU6 with 5% sucrose did not exhibit any internal levan-like structure (Fig. 1C). These data showed that the *sacB* product is a toxin in *F. nucleatum* when cells are grown in sucrose.

### Use of *sacB* as a counterselection marker for in-frame deletion in *F. nucleatum*

The ectopic expression of *sacB* inhibited fusobacterial cell growth, indicating its potential suitability for counter-selection in *F. nucleatum*. To test this, we used pCM-*galK*, a suicide plasmid with *galK* as the counterselection marker (31), to construct the plasmid pZP02 (Fig. 2A). In this plasmid, the *galK* gene and its original promoter in pCM-*galK* were replaced by *sacB* and the *rpsJ* promoter. We then generated an allelic exchange vector named pZP02-Δ*tonB*, designed to delete the entire gene region from the start to the stop codon of the *t*on*B* gene. The TonB protein, part of the TonB system along with ExbB and ExbD proteins, couples the proton motive force (PMF) of the cytoplasmic membrane to energize active transport across the outer membrane via TonB-dependent transporters (32, 33). This system is implicated in nutrient import and the secretion of intracellular proteins (34). In some bacteria, the deletion of *tonB* results in sensitivity to certain antibiotics (35, 36).

Our experiments revealed that *F. nucleatum* is sensitive to 1 µg/ml carbenicillin, and sublethal carbenicillin (0.2 µg/ml) induces the high expression of *tonB*, *exbB*, and *exbD* (unpublished data), organized in an operon (Fig. 2A). To further investigate the role of the TonB system in carbenicillin resistance, we attempted to delete the *tonB* gene using pZP02-Δ*tonB*. This plasmid contained approximately 1.5 kb of upstream and downstream DNA sequences flanking the *tonB* gene. We transformed pZP02-Δ*tonB* into *F. nucleatum* ATCC 23726 via electroporation. Selection for the in-frame deletion plasmid integration into the bacterial chromosome was performed using 5 µg/ml thiamphenicol, a chloramphenicol derivative.

We randomly picked a thiamphenicol-resistant pZP02-Δ*tonB* integrated strain. We tested the expression of *sacB* in the suicide plasmid (single copy of *sacB*), not the multiple copies of *sacB* in shuttle plasmid pZP01 (Fig. 1A), which still caused cell death in the presence of 5% sucrose. This was confirmed (see Fig. 2B). Therefore, the integration strain was continued for *sacB*/sucrose counterselection. We grew this strain overnight in the absence of antibiotics and then diluted it by 1:1000 the following day before plating cells on TSPC agar plates with 5% sucrose. To estimate the frequency of plasmid excision, a plate without sucrose was used to grow the un-counter-selected cells as a control. As shown in Figure 2C, the sucrose-negative selection only allowed a small fraction of the cells to survive. Based on the ratio of sucrose-resistant cells to total cells (cells grown on plates without selection, see Figure 2C, the left plate without sucrose), we estimated that approximately 1 in 2000 cells underwent a second recombination to lose the integrated plasmid.

Ten randomly selected sucrose-resistant colonies were tested for antibiotic sensitivity to screen for possible in-frame deletion mutants. Next, colony PCR was used to detect their genotypes using the primer pair Δ*tonB*-F/R illustrated in Figure 2A. Figure 2D shows six of the ten colonies generated amplicons of approximately 0.5 kb, indicating an in-frame deletion. The remaining four colonies produced the expected wild-type amplicons of approximately 1.5 kb. Interestingly, we found that the deletion of *tonB* indeed conferred sensitivity to carbenicillin in a dose-dependent manner (Fig. 2E) but not to erythromycin (Fig. 2F)

### Use of *sacB* as a counterselection marker for gene knockin in *F. nucleatum*

The application of counterselection markers facilitates not only markerless gene deletion but also precise gene knockin at specific sites (16). To further test the *sacB*/sucrose system in *F. nucleatum*, we attempted to insert a luciferase gene directly after the *hslO* gene, aiming to monitor the expression of the *hslO*-located gene operon in real time. The *hslO* gene encodes the zinc-dependent, redox-regulated chaperone known as Hsp33 and is the last gene of a five-gene operon (*comM-pilT-hemN-comEA-hslO*) (37) (Fig. 3A). ComM is a hexametric helicase promoting branch migration during natural transformation in Gram-negative species (38); PilT is crucial for the twitching motility of bacteria with type IV pili, aiding in bringing DNA close to the cell surface and enabling its uptake (39). HemN is an oxygen-independent coproporphyrinogen III oxidase catalyzing the conversion of coproporphyrinogen III to protoporphyrinogen IX, a critical step in the heme biosynthesis pathway under anaerobic conditions. ComEA is a membrane-associated DNA-binding protein involved in bacterial competence. ComM, PilT, and ComEA are known for their involvement in bacterial natural competence (39), suggesting this 5-gene operon is related to fusobacterial natural competence. *F. nucleatum* has demonstrated natural competence ability related to type IV pilus expression (40). However, there is no evidence yet of type IV pilus expression on the surface of *F. nucleatum*, although its genome contains an intact gene operon for type IV pilus formation (41). One possibility is that type IV pilus formation may occur in a short time window or under specific conditions. Tracing the expression of this operon may provide some answers.

**Figure 3.**
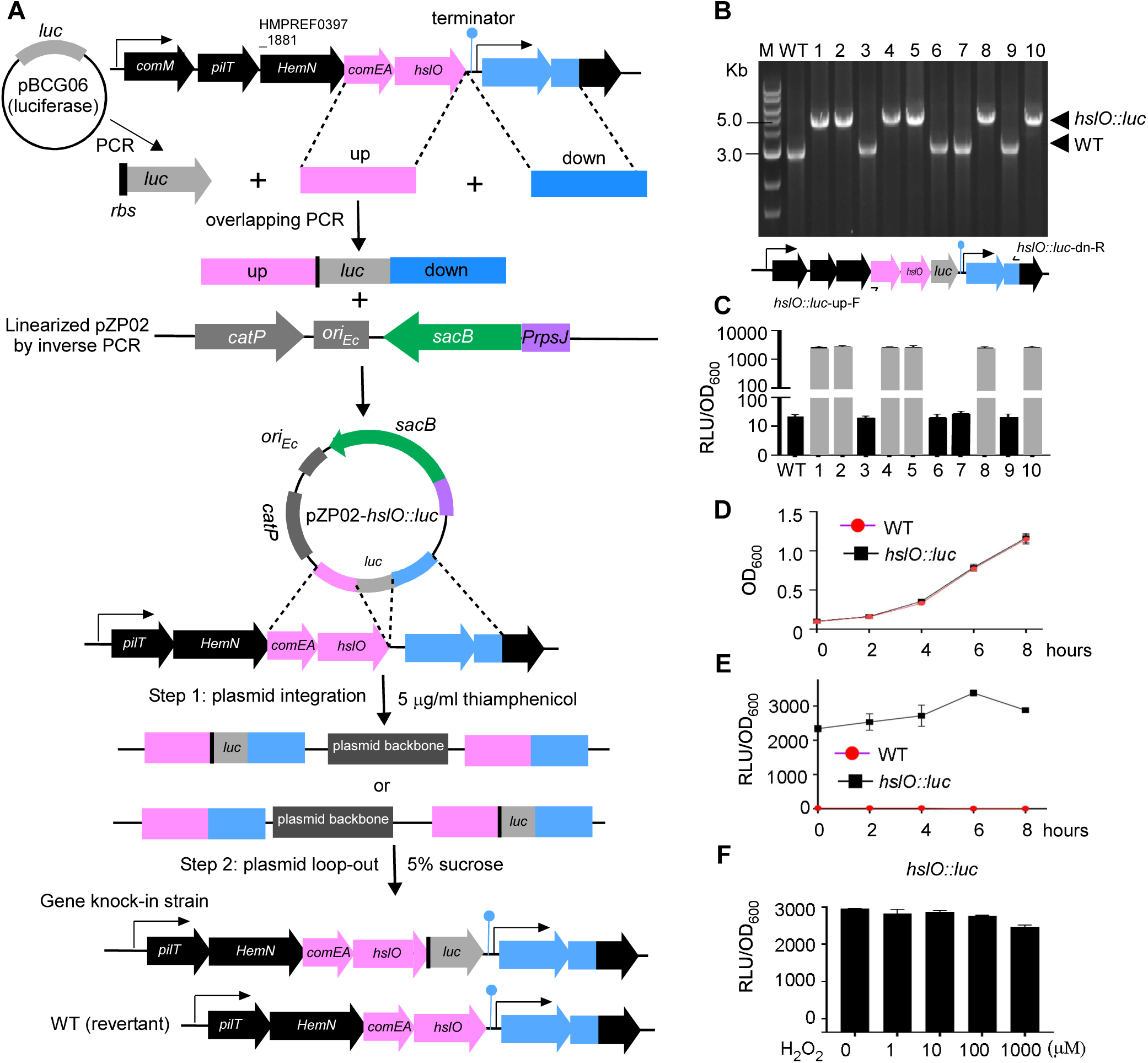
Construction of *F. nucleatum* luciferase reporter using sacB-mediated *luc* gene knock-in. (A) The *luc* gene from pBCG06 and 1.5 kb upstream and downstream regions adjacent to the *hslO* stop codon were PCR-amplified and fused via overlapping PCR. The fused product was ligated into linearized pPZ02 using Gibson assembly, creating pPZ02-*hslO::luc*. This plasmid was used for *luc* gene knock-in to generate the *hslO* luciferase reporter strain in *F. nucleatum* via a two-step allelic exchange. (B) Verification of *hslO::luc* reporter strains by colony PCR using primers *hslO::luc*-upF/*hslO::luc*-dnF. The wild-type yields a 3.0 kb amplicon, while the *hslO::luc* reporter strains yield a 4.7 kb amplicon. Six out of ten colonies were confirmed as *luc* knock-in strains. (C, D) *F. nucleatum* wild-type and *hslO::luc* strains were diluted 1:10 in fresh TSPC broth and incubated at 37°C for 8 hours. Luciferase activity and bacterial growth (OD_600)_ were measured every 2 hours, with luciferase activity normalized to OD_600_ (RLU/OD_600_). (E) The *hslO::luc* strain was grown to log phase (OD_600_ ∼0.8) and stimulated with H₂O₂ (0-1,000 µM) for 30 minutes. Luciferase activity was normalized to OD_600_ and is presented as RLU/OD_600_. Results are means ± SD from three independent experiments.

We planned to insert a DNA fragment, including the entire gene sequence of the Luciola red luciferase gene (*luc*) (42) and its RBS, immediately after the stop codon of the *hslO* gene and before the transcriptional terminator (Fig. 3A). Two DNA fragments of equal length, located before and after the *hslO* gene stop codon, were amplified via PCR and fused with *luc* between them using overlapping PCR. The fused product (up-*luc*-down) was cloned into the linearized pZP02, resulting in the plasmid pZP02-*hslO::luc*. We transformed this plasmid into the wild-type strain ATCC 23726 and selected the plasmid-integrated strains in the presence of thiamphenicol. To facilitate the excision of the integrated plasmid from the chromosome (Fig. 3A), we grew the integrated strains in the absence of antibiotics. The following day, aliquots of 1000-fold diluted cultures were plated on TSPC agar plates containing 5% sucrose. After three days, ten sucrose-resistant colonies were randomly chosen to check thiamphenicol sensitivity (100% efficiency). Colony PCR was then used to analyze the genotypes of the ten colonies using the primer pair *hslO::luc*-up-F and *hslO::luc*-dn-R (see Table 2) indicated in Figure 3B. The results revealed that six of the colonies were the expected *luc* knockin strains, while four were the wild-type revertant strains (Fig. 3B). Lastly, the *luc* knockin strains were confirmed by measuring luciferase activity (Fig 3C). The *luc* knockin at the site of *hslO* did not affect cell growth compared to the wild-type strain (Fig. 3D). When grown in a TSPC medium under anaerobic conditions, the *hslO* gene operon expression increased at the mid-log phase but subsequently decreased (Fig. 3E). Since Hsp33 is a redox-regulated chaperone that becomes activated in response to oxidative stress, such as exposure to H₂O₂ (43), we reasoned that H₂O₂ would affect this gene operon. However, adding different amounts of H₂O₂ did not change the luciferase activity in our tested conditions (Fig. 3F).

## DISCUSSION

In this study, we developed and validated a *sacB*/sucrose-based counterselection system for efficient allelic exchange in *F. nucleatum*. This method exploits the lethal sensitivity of *F. nucleatum* to sucrose when the *sacB* gene is expressed, offering a robust mechanism for creating unmarked gene deletions and precise gene knock-ins.

SacB, originally from *Bacillus subtilis*, is the bacteria’s most commonly used counterselection marker (16). We observed that the expression of SacB in *F. nucleatum* led to the accumulation of large granular particles inside the cytoplasm, causing the rupture of the inner cell membrane and resulting in cell death (Fig. 1E). This phenomenon is similar to what has been reported in *Streptococcus agalactiae*, where the expression of SacB in the presence of sucrose also leads to the formation of large intracellular inclusion bodies (30). Interestingly, in *Fusobacterium mortiferum*, the addition of serine to the culture medium in the presence of sucrose also results in the accumulation of intracellular glycogen granules(44). However, no *sacB* homolog could be identified in the genome of *F. mortiferum* or other fusobacteria. In *F. nucleatum*, the accumulation of granular sugar polymers has been observed when galactose or glucose is present, but this does not lead to cell death (24). The specific toxicity of levansucrase-produced levan in fusobacterial cells remains an intriguing question and warrants further study.

Compared to the *galK* counterselection marker we developed earlier (17), the apparent advantage of the *sacB*/sucrose system is its ability to work directly on a wild-type background strain. Previous studies have highlighted a limitation of the *sacB*/sucrose system: the occurrence of false positives due to the relatively large size of the *sacB* gene, which increases the likelihood of spontaneous mutations that render SacB nonfunctional(45, 46). However, in our experiments, we have not observed such phenomena yet. All colonies tested after *sacB*/sucrose counterselection had lost the plasmids, similar to our recently developed HicA system (47). Nonetheless, compared to the HicA counterselection system, the *sacB*/sucrose method has a drawback: adding 5% sucrose to the medium affects the growth of *F. nucleatum*. Typically, colonies under HicA selection conditions reach a pickable size (2mm in diameter) within three days, whereas in the presence of sucrose, this takes 4.5 days. Despite this limitation, the *sacB/*sucrose system complements other counterselection systems available for *F. nucleatum*, enriching the current toolkit. Given the ease of access to sucrose, we anticipate that this method will be widely adopted in the field, advancing research into *F. nucleatum* and its role in human health and disease.

## MATERIALS AND METHODS

### Bacterial strains, media, and growth conditions

Bacterial strains and plasmids used in this study are listed in Table 1. *F. nucleatum* strains were grown in Tryptic Soy Broth (TSB) supplemented with 1% Bacto^TM^ Peptone and 0.05% freshly made cysteine (TSPC) or on TSPC agar plates anaerobically (80% N_2_, 10% CO_2_, 10% H_2_) at 37°C, for the selection and counter-selection of transformants, cultures, and plates were supplemented with thiamphenicol at 5 µg/ml or 5% sucrose. *Escherichia coli* strains were grown in Luria–Bertani (LB) broth (Difco) with aeration at 37°C. *E. coli* strains carrying plasmids were grown in LB media containing 20 µg/ml chloramphenicol (Sigma).

**Table 1:** Bacterial strains and plasmids used.

| Strain & Plasmid | Description | Reference |
| --- | --- | --- |
| **Strains** |  |  |
| *F. nucleatum* 23726 | Parental strain (wild-type strain) | From ATCC |
| *F. nucleatum* ZP12 | WT strain 23726 containing pZP01 | This study |
| *F. nucleatum* ZP13 | $\Delta tonB$; an isogenic derivative of 23726 | This study |
| *F. nucleatum* ZP14 | *hsIO::luc*; an isogenic derivative of 23726 | This study |
| **Plasmids** |  |  |
| pCWU6 | *E. coli*/*Fusobacterium* shuttle vector, chloramphenicol /thiamphenicol resistance; $cm^R$/thia<sup>R</sup> | (17) |
| pCM-galk | *Clostridium perfringens* vector expressing *galk* Gene | (31) |
| pK18sB | replacement plasmid with a *SacB* as a selection marker | From Addgene |
| pZP01 | Derivative of pCWU6, mCherry gene was replaced by *P<sub>rpsJFn</sub>-sacB* | This study |
| pZP02 | Derivative of pCM-galk; *galk* gene and its promoter was replaced by *P<sub>rpsJFn</sub>-sacB* | This study |
| pZP02-$\Delta tonB$ | Derivative of pZP02; *tonB* deletion plasmid | This study |
| pZP02-*hsIO::luc* | Derivative of pZP02, transcriptional fusion *luc* gene with *hsIO* | This study |

### Plasmid construction

In this study, plasmids were created using the Gibson Assembly Master Mix kit (New England Biolabs, catalog #: E5520S) according to the manufacturer’s guidelines and were listed in Table 1. A repliQa HiFi ToughMix from Quantabio (Catalog #: 95200) was used for all PCR reactions. A linearized vector, obtained from inverted PCR or restriction enzyme digestion, was combined with inserted PCR fragments and 2x Gibson Assembly Master Mix. This mixture, with a volume of 20 µl, was incubated at 50°C for 20 minutes. Subsequently, 2 µl of the mixture was introduced into *E. coli* competent cells per the transformation protocol. To verify the generated plasmids, PCR and Sanger sequencing were carried out. The PCR primers used are listed in Table 2. All plasmids were propagated using *E. coli* DH5α as the cloning strain and later electroporated into *F. nucleatum* ATCC 23726 (48).

**Table 2:**
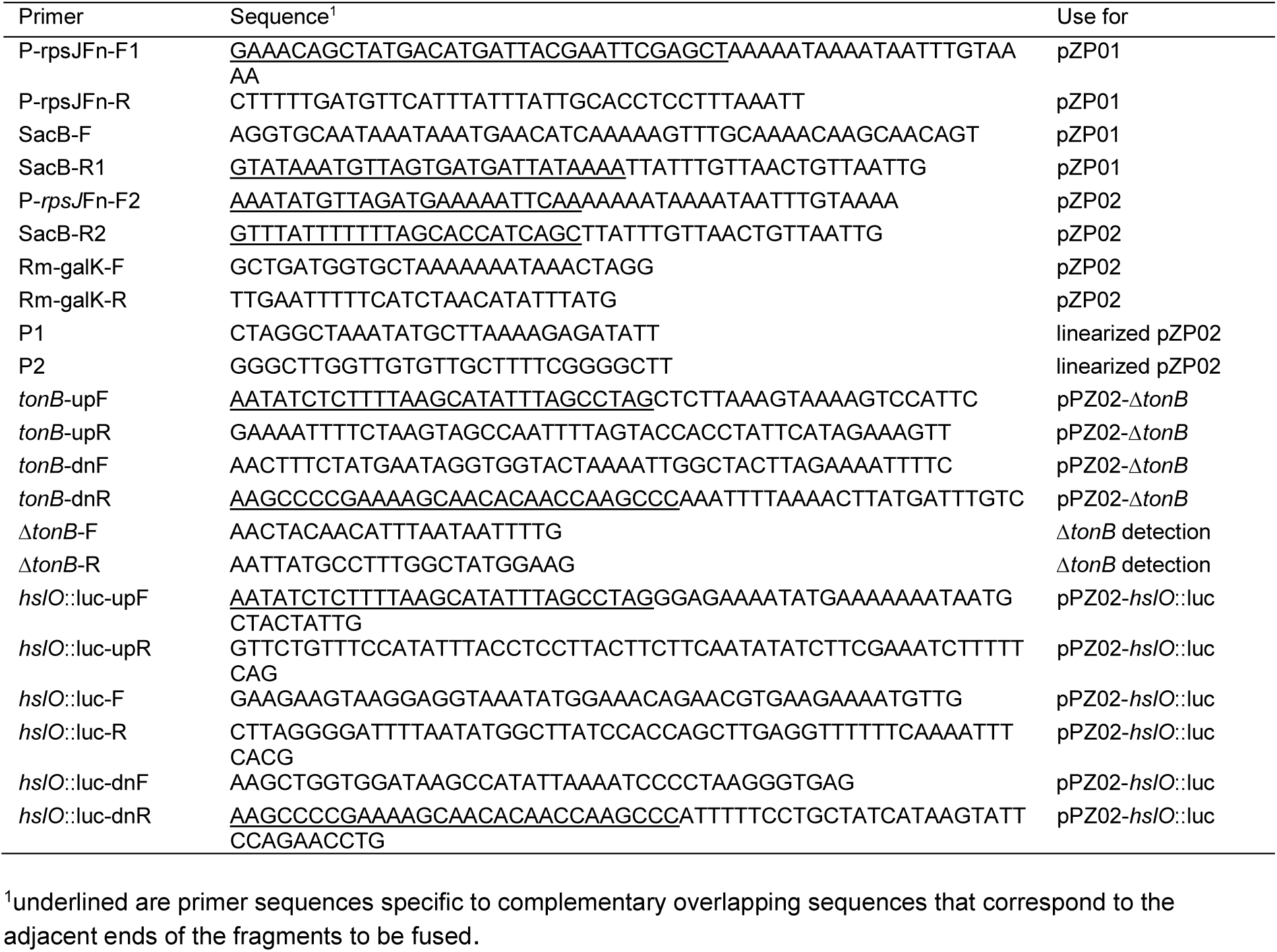
Primers used in this study.

#### (i) pZP01

To determine the sucrose sensitivity of the *F. nucleatum* strain expressing *sacB*, we created pZP01. The backbone was built using the *E. coli/F. nucleatum* shuttle vector pCWU6 (17). The primers P*rpsJ*-F1 (F stands for forward) and P*rpsJ*-R (R stands for reverse) were used to amplify the promoter region of *rpsJ* (HMPREF0397_1915; http://mg.jgi.doe.gov) from *F. nucleatum* ATCC 23726 with genomic DNA as the template. The *sacB* sequence was amplified from the pK18sB plasmid (49) obtained from Addgene using the primers *sacB*-F and *sacB*-R1. Using primer pair P*rpsJ*-F1 and *sacB*-R1, an overlapping PCR fused the two PCR products. The resultant PCR product was cloned into the SacI-and HindIII-digested pCWU6 via Gibson assembly, creating pZP01 (Fig. 1A).

#### (ii) pZP02

The pCM-galK suicide plasmid, used for in-frame gene deletion in *F. nucleatum* (17), employs the *galK* gene from *Clostridium acetobutylicum* ATCC 824 as a counterselection marker (31), controlled by the *fdx* promoter. We intend to substitute *galK* with *sacB* as a new selection marker. Using an inverse PCR with the primer pair RM-galK-F/R, the *fdx* promoter and *galK* from pCM-galK were concurrently removed, leaving a linearized pCM-galK backbone. The PrpsJ-F2 and sacB-R2 primers were used to amplify the *rpsJ* promoter and *sacB* gene from pZP01. Following this, the PCR product (*PrpsJ-sacB*) was ligated with the linearized plasmid backbone using a Gibson assembly reaction, forming the pZP02 (Fig. 2A).

#### (iii) pZP02-Δ*tonB*

To create an in-frame deletion (from the start codon to the stop codon) construction of *tonB* (HMPREF0397_0652; accessible at http://mg.jgi.doe.gov), 1.5-kb fragments upstream and downstream of *tonB* were amplified by PCR with primer pairs *tonB*-up-F/*tonB*-up-R and *tonB*-dn-F/*tonB*-dn-R, respectively. An overlapping PCR then ligated the two PCR amplicons. To clone the ligated fragment into pZP02, the pPZ02 was linearized by an inverse PCR with the primers P1 and P2 (Figure 2A). Next, a Gibson assembly reaction joined the ligated fragment and linearized the pPZ02 backbone. PCR and sequencing confirmed and validated the resulting plasmid pPZ02-Δ*tonB*.

#### (IV) pZP02-*hslO*::*luc*

The constructed plasmid is designed for the precise insertion of the luciferase gene (*luc*) immediately after *hslO* (HMPREF0397_1879; accessible at http://mg.jgi.doe.gov), which is the last gene in a five-gene operon (*comM-pilT-hemN-comEA-hslO*) (37). Specifically, our purpose is to insert the *luc* gene directly post the *hslO* stop codon, yet preceding the transcriptional terminator of this operon in the chromosome of *F. nucleatum* (Figure 3A). To do so, the primer pair *hslO::luc*-upF/R was first used to amplify a 1.5 kb fragment situated before and inclusive of the stop codon of *hslO*. The PCR product was labeled as the upstream fragment (up, see Fig. 3A). Next, a separate 1.5 kb sequence following the stop codon using primers *hslO::luc*-dnF/R was also amplified, which we referred to as the downstream fragment (down, see Figure 3). In the next step, the *luc* gene, which included a ribosome binding site (GGAGG), was amplified using pBCG06 (50) as a DNA template with primers *luc*-F/*luc*-R. The three amplicons (upstream, downstream fragments, and *luc* gene) contain overlapping regions, enabling us to carry out an overlapping PCR with the primer pair *hslO::luc*-upF and *hslO::luc*-dnR. The resultant 3.7-kb tripartite fusion product was cloned into the pZP02 plasmid, which had been linearized via an inverse PCR with the primers P1/P2 (Table 2, Fig. 3A). This was achieved using the Gibson Assembly method. We validated our construct by plasmid sequencing, confirming the successful insertion of the *luc* gene at the site we had initially intended (Fig.3A).

### Sucrose sensitivity assay

To assess if the *sacB* works as a counterselection marker, wild-type strains containing pZP01 or pCWU6 (an empty plasmid serving as a control) were initiated from a single colony and grown overnight. These two overnight cultures were subsequently streaked side-by-side onto TSPC agar plates with varying sucrose concentrations, ranging from 0 to 5%. The plates were then incubated in an anaerobic chamber at 37°C for two days before being imaged.

### Gene deletion or insertion based on sucrose counterselection in *F. nucleatum*

The plasmids pPZ02-Δ*tonB* and pPZ02-*hlsO::luc* were each introduced into *F. nucleatum* ATCC 23726 competent cells via electroporation (48). The integration of these plasmids into the chromosomal DNA through homologous recombination was selected on TSPC agar plates, which were supplemented with 5 µg/ml of thiamphenicol. Colonies that emerged on the antibiotic-laden plates were transferred and allowed to grow overnight in a TSPC broth devoid of antibiotics. The following day, these cultures were diluted by 1,000 and 10,000 in fresh TSPC media. A 100 µl sample from each dilution was then spread on TSPC agar plates containing 5% sucrose. The medium allowed us to select the cells where the integrated plasmid had been removed due to a second recombination event. After a growth period of three days in an anaerobic chamber, we chose about 20 colonies. They were individually re-streaked on both regular TSPC plates and those supplemented with 5 µg/ml of thiamphenicol to assess their sensitivity to this antibiotic. They all exhibited sensitivity to thiamphenicol, indicating a loss of the integrated plasmids. 10 of them were subjected to PCR analysis to further confirm the in-frame deletion or insertion of the target locus (Figures 2 and 3).

### Antibiotics resistance assays

To examine the *tonB* effect on antibiotic resistance, we initiated overnight cultures from a single colony of either the *F. nucleatum* wild-type strain or the Δ*tonB* mutant strain. Following the overnight culture, we centrifuged the samples to collect pellets. Each of these pellets was then individually resuspended in fresh media and diluted to an optical density at 600 nm (OD_600_) approximately equal to 0.02. We then distributed these resulting diluted cultures into a series of 8 ml TSPC medium tubes, each supplemented with varying concentrations of either erythromycin or carbenicillin. The next step involved a 2-day incubation period in an anaerobic chamber set at 37°C. Post incubation, the cultures’ OD_600_ values were measured using a spectrophotometer. The experiments were repeated three times.

### Luciferase assays

We initiated overnight cultures of these strains to study the growth and luciferase activities of F. nucleatum wild-type and the *hslO::luc* reporter strains. Post the overnight culture period, these strains were diluted in a 1:10 ratio into fresh TSPC broth. The cultures were then incubated anaerobically for 8 hours. During this period, every 2 hours, we collected three 100 µl aliquots from each culture and placed them into a 96-well microplate (#3917, Corning®). Each aliquot was then mixed with 25 μl of 1 mM D-luciferin (Molecular Probe) suspended in 100 mM citrate buffer (pH 6.0), and the mixture was vigorously pipetted 15 times under aerobic conditions. Luciferase activity was measured by using a GloMax® Navigator Microplate Luminometer. The growth of the culture was measured spectrophotometrically by determining optical density at 600 nm.

### Preparation of frozen-hydrated specimens, CryoET data collection, and 3-D reconstructions

Bacterial cells were mixed with 10 nm diameter colloidal gold beads (used as fiducial markers in image alignment) and deposited onto freshly glow-discharged, holey carbon grids for 1 min. The grids were then blotted with filter paper and rapidly frozen in liquid ethane using a gravity-driven plunger apparatus as previously described (51). The data were collected using an FEI Titan Krios transmission electron microscope, operated at 300kV and equipped with a Gatan GIF Quantum energy filter. The micrographs were recorded on a K2 Summit direct electron detector (Gatan) with a magnification of ×42,000, resulting in an effective pixel size of 3.5 Å at the specimen level. We used SerialEM (52) to collect low-dose, single-axis tilt series with dose fractionation mode at about 6 µm defocus. Tilt series were collected from -51° to 51° with increments of 3° and a cumulative dose of ∼70 e− /Å^2^ distributed over 35 stacks. Each stack contains ∼10 images. We used Tomoauto (53) to facilitate data processing, which includes drift correction of dose-fractionated data using Motioncorr (54) and assembly of corrected sums into tilt series, automatic fiducial seed model generation, alignment and contrast transfer function correction of tilt series by IMOD (55), and weighted back projection (WBP) reconstruction of tilt series into tomograms using Tomo3D (56)

## AUTHOR CONTRIBUTIONS

P.Z. and C.W. conceived and designed all experiments. P.Z., B.G., H. B., and C.W. performed all experiments. P.Z., H. B., and C. W. analyzed data. C.W. and P.Z. wrote the manuscript with contribution and approval from all authors.

## DATA AVAILABILITY STATEMENT

Materials are available upon reasonable request with a material transfer agreement with UTHealth for bacterial strains or plasmids. Plasmids information can be accessed on Benchling links at https://benchling.com/s/seq-AhrP84xr5j4gmZg5VBhc?m=slm-O4p6soWa2b9c7Rtd9Y31.

## ACKNOWLEDGMENTS

This work was supported by NIDCR grant DE030895 to C.W and NIH1R35GM138301 to B.H. We thank the UTHealth CryoEM Core Facility for providing access to its resources.

## REFERENCES

1. Paster BJ, Boches SK, Galvin JL, Ericson RE, Lau CN, Levanos VA, Sahasrabudhe A, Dewhirst FE. 2001. Bacterial diversity in human subgingival plaque. Journal of bacteriology 183:3770–3783.

2. MooreL W, Moore V. 1994. The bacteria of periodontal disease. Periodontology 2000 5:66–77.

3. Bolstad A, Jensen HB, Bakken V. 1996. Taxonomy, biology, and periodontal aspects of Fusobacterium nucleatum. Clinical microbiology reviews 9:55–71.

4. Curtis MA, Diaz PI, Van Dyke TE. 2020. The role of the microbiota in periodontal disease. Periodontology 2000 83:14–25.

5. Abusleme L, Dupuy AK, Dutzan N, Silva N, Burleson JA, Strausbaugh LD, Gamonal J, Diaz PI. 2013. The subgingival microbiome in health and periodontitis and its relationship with community biomass and inflammation. The ISME journal 7:1016–1025.

6. Han YW. 2015. Fusobacterium nucleatum: a commensal-turned pathogen. Current opinion in microbiology 23:141–147.

7. Brennan CA, Garrett WS. 2019. Fusobacterium nucleatum—symbiont, opportunist and oncobacterium. Nature Reviews Microbiology 17:156–166.

8. Kaplan C, Lux R, Huynh T, Jewett A, Shi W, Haake SK. 2005. Fusobacterium nucleatum apoptosis-inducing outer membrane protein. Journal of dental research 84:700–704.

9. Haake SK, Yoder S, Gerardo SH. 2006. Efficient gene transfer and targeted mutagenesis in Fusobacterium nucleatum. Plasmid 55:27–38.

10. Han YW, Ikegami A, Chung P, Zhang L, Deng CX. 2007. Sonoporation is an efficient tool for intracellular fluorescent dextran delivery and one-step double-crossover mutant construction in Fusobacterium nucleatum. Applied and environmental microbiology 73:3677–3683.

11. Han YW, Ikegami A, Rajanna C, Kawsar HI, Zhou Y, Li M, Sojar HT, Genco RJ, Kuramitsu HK, Deng CX. 2005. Identification and characterization of a novel adhesin unique to oral fusobacteria. Journal of bacteriology 187:5330–5340.

12. Kaplan CW, Lux R, Haake SK, Shi W. 2009. The Fusobacterium nucleatum outer membrane protein RadD is an arginine-inhibitable adhesin required for inter-species adherence and the structured architecture of multispecies biofilm. Molecular microbiology 71:35–47.

13. Lima BP, Shi W, Lux R. 2017. Identification and characterization of a novel Fusobacterium nucleatum adhesin involved in physical interaction and biofilm formation with Streptococcus gordonii. Microbiologyopen 6:e00444.

14. Casasanta MA, Yoo CC, Smith HB, Duncan AJ, Cochrane K, Varano AC, Allen-Vercoe E, Slade DJ. 2017. A chemical and biological toolbox for Type Vd secretion: Characterization of the phospholipase A1 autotransporter FplA from Fusobacterium nucleatum. Journal of Biological Chemistry 292:20240–20254.

15. Bedree JK, Bor B, Cen L, Edlund A, Lux R, McLean JS, Shi W, He X. 2018. Quorum sensing modulates the epibiotic-parasitic relationship between Actinomyces odontolyticus and its Saccharibacteria epibiont, a Nanosynbacter lyticus strain, TM7x. Frontiers in microbiology 9:2049.

16. Reyrat J-M, Pelicic V, Gicquel B, Rappuoli R. 1998. Counterselectable markers: untapped tools for bacterial genetics and pathogenesis. Infection and immunity 66:4011–4017.

17. Wu C, Al Mamun AAM, Luong TT, Hu B, Gu J, Lee JH, D’Amore M, Das A, Ton-That H. 2018. Forward Genetic Dissection of Biofilm Development by Fusobacterium nucleatum: Novel Functions of Cell Division Proteins FtsX and EnvC. mBio 9.

18. Wu C, Chen YW, Scheible M, Chang C, Wittchen M, Lee JH, Luong TT, Tiner BL, Tauch A, Das A, Ton-That H. 2021. Genetic and molecular determinants of polymicrobial interactions in Fusobacterium nucleatum. Proc Natl Acad Sci U S A 118.

19. Casasanta MA, Yoo CC, Udayasuryan B, Sanders BE, Umaña A, Zhang Y, Peng H, Duncan AJ, Wang Y, Li L. 2020. Fusobacterium nucleatum host-cell binding and invasion induces IL-8 and CXCL1 secretion that drives colorectal cancer cell migration. Science signaling 13:eaba9157.

20. Umaña A, Nguyen TT, Sanders BE, Williams KJ, Wozniak B, Slade DJ. 2022. Enhanced Fusobacterium nucleatum Genetics Using Host DNA Methyltransferases To Bypass Restriction-Modification Systems. Journal of Bacteriology 204:e00279–22.

21. Ryu E-J, Sim J, Sim J, Lee J, Choi B-K. 2016. D-Galactose as an autoinducer 2 inhibitor to control the biofilm formation of periodontopathogens. Journal of Microbiology 54:632–637.

22. Abed J, Emgård JE, Zamir G, Faroja M, Almogy G, Grenov A, Sol A, Naor R, Pikarsky E, Atlan KA. 2016. Fap2 mediates Fusobacterium nucleatum colorectal adenocarcinoma enrichment by binding to tumor-expressed Gal-GalNAc. Cell host & microbe 20:215–225.

23. Parhi L, Abed J, Shhadeh A, Alon-Maimon T, Udi S, Ben-Arye SL, Tam J, Parnas O, Padler-Karavani V, Goldman-Wohl D, Yagel S, Mandelboim O, Bachrach G. 2022. Placental colonization by Fusobacterium nucleatum is mediated by binding of the Fap2 lectin to placentally displayed Gal-GalNAc. Cell Rep 38:110537.

24. Robrish SA, Oliver C, Thompson J. 1987. Amino acid-dependent transport of sugars by Fusobacterium nucleatum ATCC 10953. Journal of bacteriology 169:3891–3897.

25. Vaillancourt K, Bédard N, Bart C, Tessier M, Robitaille G, Turgeon N, Frenette M, Moineau S, Vadeboncoeur C. 2008. Role of galK and galM in galactose metabolism by Streptococcus thermophilus. Applied and environmental microbiology 74:1264–1267.

26. Neves AR, Pool WA, Solopova A, Kok J, Santos H, Kuipers OP. 2010. Towards enhanced galactose utilization by Lactococcus lactis. Applied and environmental microbiology 76:7048–7060.

27. Chambert R, Petit-Glatron M-F. 1989. Study of the effect of organic solvents on the synthesis of levan and the hydrolysis of sucrose by Bacillus subtilis levansucrase. Carbohydrate research 191:117–123.

28. Blomfield I, Vaughn V, Rest R, Eisenstein BI. 1991. Allelic exchange in Escherichia coli using the Bacillus subtilis sacB gene and a temperature-sensitive pSC101 replicon. Molecular microbiology 5:1447–1457.

29. Jäger W, Schäfer A, Pühler A, Labes G, Wohlleben W. 1992. Expression of the Bacillus subtilis sacB gene leads to sucrose sensitivity in the gram-positive bacterium Corynebacterium glutamicum but not in Streptomyces lividans. Journal of bacteriology 174:5462–5465.

30. Hooven TA, Bonakdar M, Chamby AB, Ratner AJ. 2019. A counterselectable sucrose sensitivity marker permits efficient and flexible mutagenesis in Streptococcus agalactiae. Applied and environmental microbiology 85:e03009–18.

31. Nariya H, Miyata S, Suzuki M, Tamai E, Okabe A. 2011. Development and application of a method for counterselectable in-frame deletion in Clostridium perfringens. Applied and environmental microbiology 77:1375–1382.

32. Noinaj N, Guillier M, Barnard TJ, Buchanan SK. 2010. TonB-dependent transporters: regulation, structure, and function. Annual review of microbiology 64:43–60.

33. Postle K, Kadner RJ. 2003. Touch and go: tying TonB to transport. Molecular microbiology 49:869–882.

34. Saxena D, Maitra R, Bormon R, Czekanska M, Meiers J, Titz A, Verma S, Chopra S. 2023. Tackling the outer membrane: facilitating compound entry into Gram-negative bacterial pathogens. npj Antimicrobials and Resistance 1:17.

35. Calvopiña K, Dulyayangkul P, Heesom KJ, Avison MB. 2020. TonB-dependent uptake of β-lactam antibiotics in the opportunistic human pathogen Stenotrophomonas maltophilia. Molecular microbiology 113:492–503.

36. Dong Y, Li Q, Geng J, Cao Q, Zhao D, Jiang M, Li S, Lu C, Liu Y. 2021. The TonB system in Aeromonas hydrophila NJ-35 is essential for MacA2B2 efflux pump-mediated macrolide resistance. Veterinary research 52:63.

37. Ponath F, Tawk C, Zhu Y, Barquist L, Faber F, Vogel J. 2021. RNA landscape of the emerging cancer-associated microbe Fusobacterium nucleatum. Nat Microbiol 6:1007–1020.

38. Nero TM, Dalia TN, Wang JC-Y, Kysela DT, Bochman ML, Dalia AB. 2018. ComM is a hexameric helicase that promotes branch migration during natural transformation in diverse Gram-negative species. Nucleic acids research 46:6099–6111.

39. Craig L, Forest KT, Maier B. 2019. Type IV pili: dynamics, biophysics and functional consequences. Nature reviews microbiology 17:429–440.

40. Sanders BE, Umaña A, Nguyen TT, Williams KJ, Yoo CC, Casasanta MA, Wozniak B, Slade DJ. 2023. Type IV pili facilitated natural competence in Fusobacterium nucleatum. Anaerobe 82:102760.

41. Desvaux M, Khan A, Beatson SA, Scott-Tucker A, Henderson IR. 2005. Protein secretion systems in Fusobacterium nucleatum: genomic identification of Type 4 piliation and complete Type V pathways brings new insight into mechanisms of pathogenesis. Biochimica et Biophysica Acta (BBA)-Biomembranes 1713:92–112.

42. Merritt J, Senpuku H, Kreth J. 2016. Let there be bioluminescence: development of a biophotonic imaging platform for in situ analyses of oral biofilms in animal models. Environmental microbiology 18:174–190.

43. Kaidow A, Ishii N, Suzuki S, Shiina T, Endoh K, Murakami Y, Kasahara H. 2022. HslO ameliorates arrested ΔrecA polA cell growth and reduces DNA damage and oxidative stress responses. Scientific Reports 12:22182.

44. Robrish SA, Oliver C, Thompson J. 1991. Sugar metabolism by fusobacteria: regulation of transport, phosphorylation, and polymer formation by Fusobacterium mortiferum ATCC 25557. Infection and immunity 59:4547–4554.

45. Hashimoto JG, Stevenson BS, Schmidt TM. 2003. Rates and consequences of recombination between rRNA operons. Journal of bacteriology 185:966–972.

46. Wang T, Li Y, Li J, Zhang D, Cai N, Zhao G, Ma H, Shang C, Ma Q, Xu Q. 2019. An update of the suicide plasmid-mediated genome editing system in Corynebacterium glutamicum. Microbial biotechnology 12:907–919.

47. Bibek G, Zhou P, Wu C. 2023. A HicA toxin-based counter-selection marker for allelic exchange mutations in Fusobacterium nucleatum. bioRxiv doi:10.1101/2023.01.20.524997:2023.01.20.524997.

48. Peluso EA, Scheible M, Ton-That H, Wu C. 2020. Genetic Manipulation and Virulence Assessment of Fusobacterium nucleatum. Curr Protoc Microbiol 57:e104.

49. Jayakody LN, Johnson CW, Whitham JM, Giannone RJ, Black BA, Cleveland NS, Klingeman DM, Michener WE, Olstad JL, Vardon DR. 2018. Thermochemical wastewater valorization via enhanced microbial toxicity tolerance. Energy & Environmental Science 11:1625–1638.

50. Gc B, Zhou P, Wu C. 2023. HicA Toxin-Based Counterselection Marker for Allelic Exchange Mutations in Fusobacterium nucleatum. Applied and Environmental Microbiology:e00091–23.

51. Hu B, Khara P, Christie PJ. 2019. Structural bases for F plasmid conjugation and F pilus biogenesis in Escherichia coli. Proceedings of the National Academy of Sciences 116:14222–14227.

52. Mastronarde DN. 2005. Automated electron microscope tomography using robust prediction of specimen movements. Journal of structural biology 152:36–51.

53. Morado DR, Hu B, Liu J. 2016. Using Tomoauto: a protocol for high-throughput automated cryo-electron tomography. Journal of visualized experiments: JoVE.

54. Li X, Mooney P, Zheng S, Booth CR, Braunfeld MB, Gubbens S, Agard DA, Cheng Y. 2013. Electron counting and beam-induced motion correction enable near-atomic-resolution single-particle cryo-EM. Nature methods 10:584–590.

55. Kremer JR, Mastronarde DN, McIntosh JR. 1996. Computer visualization of three-dimensional image data using IMOD. Journal of structural biology 116:71–76.

56. Agulleiro J-I, Fernandez J-J. 2015. Tomo3D 2.0–exploitation of advanced vector extensions (AVX) for 3D reconstruction. Journal of structural biology 189:147–152.

57. Gur C, Ibrahim Y, Isaacson B, Yamin R, Abed J, Gamliel M, Enk J, Bar-On Y, Stanietsky-Kaynan N, Coppenhagen-Glazer S. 2015. Binding of the Fap2 protein of Fusobacterium nucleatum to human inhibitory receptor TIGIT protects tumors from immune cell attack. Immunity 42:344–355.

